## Supplementary tables, figures. discussion. for "Tyrosine 121 moves revealing a druggable pocket that couples catalysis to ATP-binding in serine racemase"

### Supporting information content:

**Supplementary Table 1.** Data collection and refinement statistics for seven human SR crystal structures.
**Supplementary Table 2.** Twelve XChem compounds with electron density in at least one Tyr 121 pocket and % inhibition hSR data at 10mM and 1mM.
**Supplementary Table 3.** Phi-psi angles for the β3-α4 loop and for the β6 strand regions**.**
**Supplementary Table 4.** SR XChem fragment hit information.
**Supplementary Table 5**. List of serine racemase (SR) and serine dehydratase (SDH) structures from the PDB.
**Supplementary Table 6**. Summary of SR inhibitors and physiological modulators

**Supplementary Figure 1**. Chemical structures of PLP, PLP-Lys56 and PLP-L-Ser, PLP-D-Ser and PLP-planar intermediate
**Supplementary Figure 2**. Comparison of ‘open’ and ‘closed’ structures of human and yeast serine racemase
**Supplementary Figure 3**. Serine racemase chemistry and lysino-d-alanyl and gem-diamine structures.
**Supplementary Figure 4**. Malonate binding sites in human SR crystal structures (6ZSP and 3L6B)
**Supplementary Figure 5**. Tyr121 can form hydrogen bonds to ATP or main-chain carbonyl of Ser84
**Supplementary Figure 6**. The side-chain of Cys 113 is accessible in the ‘open’ human SR structure but is buried in the ‘closed’ structure
**Supplementary Figure 7**. Serine racemase sequence alignment
**Supplementary Figure 8**. Superpositions of human *holo* and ATP/malonate-bound SR with yeast orthologues
**Supplementary Figure 9**. Superpositions of closed SR ATP/malonate-bound structures with open SR subunits A and C
**Supplementary Figure 10**. The hydrophobic core of the dimer interface in open and closed hSR structures
**Supplementary Figure 11**. Serine dehydratase and serine racemase sequence alignment
**Supplementary Figure 12**. Roles of lysine 56 and serine 84 in proton exchange in human serine racemase
**Supplementary Figure 13**. Does Arginine 135 protonate Ser 84 via the side-chain of the substrate serine?
**Supplementary Figure 14.** OMIT maps of ATP, malonate, and bound XChem ligands
**Supplementary Figure 15.** OMIT maps of XChem ligands bound to SR

**Supplementary Discussion.** The evolution and regulation of the activities of human serine racemase

**Supplementary References**

**Supplementary Table 1.** Data collection and refinement statistics for seven human SR crystal structures (including five XCHem soaks).

| Parameter/ ligand sites | Active site/ ATP | No ligand XChem control | Small domain Y121 | Sd. Y121 + L. dom. D238 | Small domain T81 | Dimer interface W278 | Dimer interface L25 |
|---|---|---|---|---|---|---|---|
| PDB ID (resolution) LIGAND | 6ZSP (1.60 Å) Mal/ATP | 6ZUJ (1.80 Å) | 7NBG (1.53 Å) 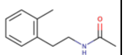 | 7NBH (1.77 Å) 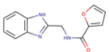 | 7NBC (1.71 Å) 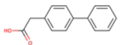 | 7NBD (1.86 Å) 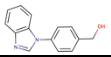 | 7NBF (1.60 Å) 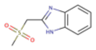 |
| Space group | $P2_12_12_1$ | $P2_1$ | $P2_1$ | $P2_1$ | $P2_1$ | $P2_1$ | $P2_1$ |
| a (Å), b (Å), c (Å) | 53.94, 81.19, 134.46 | 48.06, 155.00, 85.65 | 48.11, 154.76, 85.40 | 48.180 154.880 85.480 | 48.150 154.860 85.480 | 48.012 154.846 85.451 | 48.112 154.937 85.539 |
| $\alpha$=90°, $\beta$, $\gamma$=90° | 90° | 98.3° | 98.14° | 98.04° | 98.08° | 98.04° | 97.96° |
| Data collection parameters | | | | | | | |
| Wavelength | 0.978 | 0.91587 | 0.91587 | 0.91587 | 0.91587 | 0.91587 | 0.91587 |
| Resolution range high res. shell. | 50.1–1.60 1.63–1.60 | 84.7–1.80 1.83–1.80 | 77.4–1.53 1.57–1.53 | 51.6–1.77 1.82–1.77 | 77.4–1.71 1.74–1.71 | 84.6–1.86 1.97–1.86 | 84.7–1.60 1.63–1.60 |
| Total no. of reflections | 304094 (7908) | 381617 (19294) | 575680 (20850) | 402122 (18838) | 440513 (18933) | 333133 (42883) | 517542 (20057) |
| No. of unique reflections | 76688 (3151) | 111815 (5444) | 177718 (11899) | 116780 (8546) | 130785 (6420) | 96340 (12195) | 158956 (7536) |
| Completeness (%) | 97.5 (81.0) | 97.8 (95.61) | 95.9 (87.0) | 97.1 (96.3) | 98.2 (97.2) | 94.0 (82.0) | 97.6 (93.4) |
| Multiplicity | 4.0 (2.5) | 3.4 (3.5) | 3.2 (2.6) | 3.4 (3.3) | 3.4 (2.9) | 3.5 (3.5) | 3.3 (2.7) |
| I/σI | 9.2 (1.3) | 9.5 (1.7) | 9.0 (1.0) | 8.2 (1.0) | 8.3 (1.0) | 8.9 (1.1) | 8.9 (1.1) |
| $R_{merge}$ (I) | 0.069 (0.577) | 0.068 (0.703) | 0.052 (0.804) | 0.072 (1.048) | 0.067 (0.948) | 0.073 (0.912) | 0.059 (0.658) |
| CC1/2 | 0.999 (0.512) | 0.9973 (0.651) | 0.998 (0.453) | 0.997 (0.302) | 0.997 (0.305) | 0.998 (0.470) | 0.997 (0.383) |
| R-meas | 0.079 (0.705) | 0.081 (0.829) | 0.075 (1.002) | 0.098 (1.424) | 0.093 (1.306) | 0.100 (1.235) | 0.082 (0.889) |
| Refinement res. range (Å) | 20–1.6 | 84.1–1.8 | 77.4–1.53 | 47.7–1.77 | 77.6–1.71 | 84.6–1.86 | 84.7–1.60 |
| No. of refls, working (test) | 72966 (3722) | 106173 (5512) | 168961 (8706) | 110878 (5859) | 124213 (6527) | 91467 (4830) | 150883 (7962) |
| Final Rcryst | 0.192 | 0.179 | 0.183 | 0.172 | 0.174 | 0.176 | 0.172 |
| Final Rfree | 0.227 | 0.220 | 0.201 | 0.202 | 0.201 | 0.207 | 0.198 |
| No. of non-H atoms | | | | | | | |
| Protein | 5267 | 9879 | 10369 | 10201 | 10079 | 10155 | 10106 |
| Ions | 4 | 11 | 15 | 16 | 12 | 10 | 14 |
| Ligand/other | 125 | 202 | 239 | 294 | 184 | 245 | 284 |
| Water | 333 | 974 | 946 | 989 | 867 | 1069 | 929 |
| Total | 5729 | 11066 | 11570 | 11500 | 11142 | 11479 | 10503 |
| R.M.S. deviations | | | | | | | |
| Bonds (Å) | 0.006 | 0.006 | 0.004 | 0.009 | 0.007 | 0.010 | 0.013 |
| Angles (°) | 1.348 | 1.319 | 1.151 | 1.446 | 1.356 | 1.669 | 1.882 |
| Average B factor (Å$^2$) | 21.9 | 34.2 | 28.6 | 35.2 | 32.4 | 37.9 | 29.0 |
| Protein | 21.7 | 34.4 | 26.7 | 34.4 | 32.2 | 37.3 | 28.7 |
| Ion | 15.1 | 36.4 | 40.3 | 47.8 | 39.3 | 41.2 | 31.6 |
| Ligand | 22.7 | 38.7 | 34.8 | 44.6 | 35.5 | 39.9 | 28.4 |
| Water | 32.3 | 42.1 | 39.4 | 45.8 | 41.5 | 49.7 | 38.6 |
| Ramachandran plot | | | | | | | |
| Favoured regions (%) | 98.2 | 97.8 | 98.0 | 98.1 | 96.3 | 97.4 | 97.7 |
| Additionally allowed (%) | 1.8 | 1.1 | 1.8 | 1.8 | 3.5 | 2.5 | 2.3 |
| Outliers (%) | 0.0 | 1.1 | 0.2 | 0.1 | 0.2 | 0.1 | 0.0 |

**Supplementary Table 2.** Twelve XChem compounds with electron density in at least one Tyr 121 pocket and % inhibition hSR data at 10mM and 1mM.

| PDB code Cmp no. (Resol.) | SMILES | Picture | Initial rank*/ Comments | % Inhibition at | |
|---|---|---|---|---|---|
| | | | | 10mM | 1mM |
| 7NBG **1**-x0495 (1.53Å) | CC(=O)NCCC=1C=CC=CC1C | 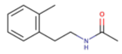 | A. Methylbenzene in pocket in subunits A and B (and half occupancy in D. Tyr 121 in pocket in C. | 102 | 33 |
| 7NBH **2**-x0406 (1.77Å) | O=C(NCC1=NC=2C=CC=CC2N1)C3=CC=CO3 | 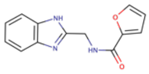 | B. Benzimidazole in subunits A,B,D. Tyr 121 in pocket in subunit C. Second site. | ND | ND |
| **3**-x0458 (1.85Å) | NC=1C=CC=CC1N2CCOCC2 | 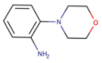 | B. Aniline in pocket in A,B subunits. Aniline nitrogen H-bonds to MC, C=0 of Ser 84 and Gly 85.? | 103 | 82 |
| **4** (x0482) (1.71Å) | CC(=O)NC1CNC=2C=CC=CC2C1 | 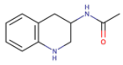 | B. Benzene in pocket in subunits A and B, nitrogen of tetrahydroquinoline may make H-bond to Ser 84 C=O. | 104 | 59 |
| **5** (x0501) (1.82Å) | NC(=O)C=1C=NC(=NC1)C(F)(F)F | 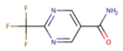 | D. Tyrosine 121 in pocket in subunits A and B, electron density in subunits C and D suggests half occupancy compound with carboxamide in pocket mimicking hydroxyl of Tyr 121? | 52 | 9 |
| **6** (x0513) (1.45Å) | CNCC1COC=2C=CC=CC2O1 | 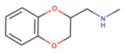 | A. Benzene in pocket in subunits A,B,D. Tyr 121 in pocket in subunit C. | 101 | 108 |
| **7** (x0109) (1.73Å) | O=C(CCCC=1C=CC=CC1)N2CCOCC2 | 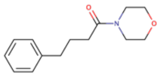 | C. Benzene in pocket in subunits A,B,D. Tyr 121 in pocket in subunit C. | ND | ND |
| **8** (x0127) (1.87Å) | CCCC1=NN=C(NC(=O)C=2C=CC=CN2)S1 | 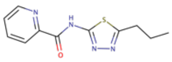 | D. Pyridine in pocket in subunit D. | ND | ND |
| **9** (x0154) (1.83Å). | CN1N=CC=C1C(=O)NCC2=CC=CS2 | 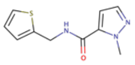 | D. Density in pocket for one end of cmpnd in subunit B and D. | ND | ND |
| **10** (x0302) (2.23Å) | CC(CS(=O)(=O)N)C=1C=CC=CC1 | 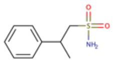 | D. Benzene in pocket in subunit A. | 64 | 29 |
| **11** (x0431) (1.51Å) | CS(=O)(=O)CC(O)C=1C=CC=CC1 | 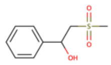 | B. Benzene in pocket in subunits A and B. | ND | ND |
| **12** (x0437) (1.74Å) | NCC1=CC(=NO1)C=2C=CC=CC2 | 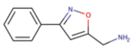 | D. Benzene in pocket in subunits A,B,D. Tyr 121 in pocket in subunit C. | ND | ND |

* Initial rank from examination of XChem electron density/fit; A = best, D = worst.

**Supplementary Table 3.** Phi-psi angles for the b3-a4 loop and for the b6 strand regions**.**

| | Holoenzyme (6ZUJ). | | | | Malonate/ATP (6ZSP) | | | |
|---|---|---|---|---|---|---|---|---|
| | A-subunit | | C-subunit | | A-subunit | | B-subunit | |
| | phi | psi | phi | psi | phi | psi | phi | psi |
| THR 81 | -153 | 157 | -138 | 148 | -158 | 178 | -157 | 176 |
| **HIS 82** | -92 | -9 | -89 | -27 | -117 | 150 | -117 | 148 |
| SER 83 | -97 | 126 | -78 | 131 | -162 | 146 | -163 | 145 |
| SER 84 | -107 | 0 | -102 | 3 | -76 | -11 | -75 | -10 |
| **GLY 85** | -97 | -128 | -92 | -153 | 150 | -45 | 150 | -44 |
| ASN 86 | -57 | -34 | -60 | -41 | -71 | -47 | -70 | -49 |
| HIS 87 | -71 | -45 | -61 | -50 | -61 | -38 | -62 | -37 |
| GLY 148 | -94 | 176 | -97 | -178 | -84 | -175 | -85 | -175 |
| ILE 149 | -109 | 136 | -123 | 156 | -113 | 123 | -112 | 129 |
| MET 150 | -88 | 129 | -116 | 148 | -81 | 130 | -82 | 127 |
| VAL 151 | -121 | 103 | -132 | 117 | -124 | 98 | -119 | 96 |

**Supplementary Table 4. Compound inhibition at 1 and 10mM.** Human SR inhibition data (see methods for details) is shown for malonate and some of the fifteen fragments from the XChem screen. ND = No data.

| Fragment number | Name or DSiP number | Abbreviation | Structure | % Inhibition at | |
|---|---|---|---|---|---|
| | | | | 10mM | 1mM |
| **1** | Z52314092 | x0495 | 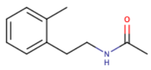 | 102 | 33 |
| **2** | Z26781964 | x0406 | 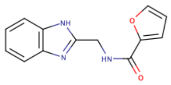 | ND | ND |
| **3** | Z56785490 | x0458 | 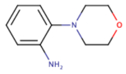 | 103 | 82 |
| **4** | Z1492796719 | x0482 | 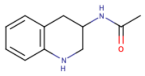 | 104 | 59 |
| **5** | Z1745658474 | x0501 | 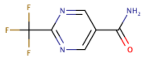 | 52 | 9 |
| **6** | Z1891773476 | x0513 | 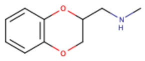 | 101 | 108 |
| **7** | Z419995480 | x0109 | 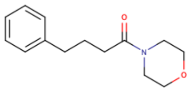 | ND | ND |
| **8** | Z52425517 | x0127 | 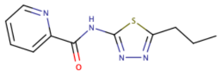 | ND | ND |
| **9** | Z915492990 | x0154 | 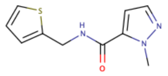 | ND | ND |
| **10** | Z1407673036 | x0302 | 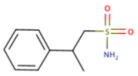 | 64 | 29 |
| **11** | Z822382694 | x0431 | 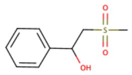 | ND | ND |
| **12** | Z425449682 | x0437 | 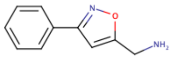 | ND | ND |
| **13** | Z2856434779 | x0430 | 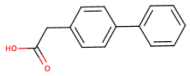 | ND | ND |
| **14** | Z235449082 | x0478 | 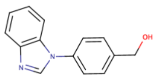 | ND | ND |
| **15** | Z126932614 | x0306 | 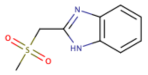 | ND | ND |

**Supplementary Table 5**. List of serine racemase (SR) and serine dehydratase (SDH) structures from the PDB

| PDB code (resolution) | Species | Protein | Ligands | Metal ion site metal | Reference |
|---|---|---|---|---|---|
| **6ZSP** (1.60 Å) | *H. sapiens* | SR | PLP, MAL, ATP | $Mg^{2+}$ | Current study |
| **6ZUJ** (1.80 Å) | | | PLP | $Ca^{2+}$ | Current study |
| **7NBC** (1.71 Å) | | | PLP, x0430 | $Ca^{2+}$ | Current study |
| **7NBD** (1.86 Å) | | | PLP, x0478 | $Ca^{2+}$ | Current study |
| **7NBF** (1.60 Å) | | | PLP, x0306 | $Ca^{2+}$ | Current study |
| **7NBG** (1.53 Å) | | | PLP, x0495 | $Ca^{2+}$ | Current study |
| **7NBH** (1.77 Å) | | | PLP, x0406 | $Ca^{2+}$ | Current study |
| **6SLH** (1.89 Å) | | | PLP | $Mg^{2+}$ | 1 |
| **5X2L** (1.81 Å) | | | PLP | $Mg^{2+}$ | 2 |
| **3L6B** (1.50 Å) | | | PLP, MAL | $Mn^{2+}$ | 3 |
| **3L6C** (2.20 Å) | *R. norvegicus* | SR | PLP, MAL | $Mn^{2+}$ | 3 |
| **3HMK** (2.10 Å) | | | PLP | $Mn^{2+}$ | 3 |
| **1WTC** (1.90 Å) | *S. pombe* | SR | PLP, AMPPCP | $Mg^{2+}$ | 4 |
| **1V71** (1.70 Å) | | | PLP | $Mg^{2+}$ | 4 |
| **2ZR8** (2.20 Å) | | | PDD, L-SER | $Mg^{2+}$ | 4 |
| **2ZPU** (1.70 Å) | | | PDD | $Mg^{2+}$ | 5 |
| **5CVC** (2.09 Å) | *Z. mays* | SR | PLP | $Mg^{2+}$ | 6 |
| **1PWE** (2.80 Å) | *R norvegicus* | SDH | - | - | 7 |
| **1PWH** (2.60 Å) | | | PLV | $K^{+}$ | 7 |
| **1P5J** (2.50 Å) | *H. sapiens* | SDH | PLP | $H_2O$ | 8 |
| **4H27** (1.30 Å) | | | PLP | $H_2O$ | 9 |
| **2RKB** (2.80 Å) | | SDH like-1 | PLP | $K^{+}$ | 10 |

AMPPCP, Adenosine 5′-[β,γ-methylene]triphosphate (non-hydrolysable analogue of ATP); ATP, Adenosine triphosphate; L-SER, L-Serine; MAL, malonate; PDD, N-(5′-phosphopyridoxyl)-D-alanine; PLP, pyridoxal 5′-phosphate; PLV, PLP-O-methylserine aldimine

**Supplementary Table 6**. Summary of SR inhibitors and physiological modulators

| Group | Compound | Inhibition mode | Reference |
|---|---|---|---|
| **Serine analogues** | L-serine-*O*-sulfate | Competitive/ substrate mimetic | 11-13 |
| | L-cysteine-*S*-sulfate | | 11 |
| | β-chloro-L-alanine | | 11,13 |
| | Homocysteic acid | | 11,13 |
| **Aspartate analogues** | L-aspartate | Competitive | 11 |
| | L-asparagine | | 11 |
| | L-erythro-3-hydroxyaspartate | | 11 |
| | L-aspartate β-hydroxamate | | 14 |
| **Carboxylic acids** | Malonic | Competitive | 11 |
| | Succinic | | 11 |
| | Maleic | | 11 |
| | L-malic | | 11 |
| | Tartaric | | 11 |
| | Dihydroxyfumaric | | 11 |
| | Oxaloacetic acid | Product inhibition | 11,13 |
| **Carboxylate-containing compounds** | Various; see reference | Competitive | 15 |
| **Malonate derivatives** | 2-aminomalonate | Competitive | 16 |
| | 2-fluoromalonate | | 16 |
| | 2-hydroxymalonate | | 16,17 |
| | 2,2-dichloromalonate | | 17 |
| | 2,2-bis(hydroxymethyl)malonate | | 17 |
| **Substrate-product analogues** | α-(hydroxymethyl)serine | Substrate-product inhibition | 18 |
| **Cyclopropane carboxylic acid derivatives** | 1,2-cyclopropanedicarboxylic acid | Competitive; related to malonate | 19 |
| **Peptidomimetic inhibitors** | Various; see references | Malonate site/ dimer interface | 2,20-22 |
| **Plant derivatives** | Madecassoside | Competitive | 23 |
| **Non-specific inhibitors** | Sulfhydryl compounds (e.g. L-cysteine) | PLP modification | 13 |
| | Hydroxamic acids | | 14 |
| **Physiological modulators** | PIP2 | Inhibition (noncompetitive); Interferes with ATP binding | 24 |
| | GRIP | Activation; binds to SR | 25 |
| | PICK1 | Inhibition; binds to SR and induces its PPN by PKC | 26 |
| | Golga3 | Increases SR levels by preventing ubiquitylation | 27 |
| | Nitric oxide | Inhibition; induces S-nitrosylation of SR | 28 |
| | NADH[9] + analogues | Inhibition; promotes ATP dissociation | 29,30 |
| | Stargazin/PSD-95 | Inhibition; facilitates cell membrane localization of SR | 31 |
| | DISC1 | Stabilises SR and prevents its degradation | 32 |

ATP, adenosine triphosphate; DISC1, Disrupted-In-Schizophrenia-1; Golga3, Golgin subfamily A member 3; GRIP, glutamate receptor-interacting protein; NADH, nicotinamide adenine dinucleotide; PICK1, protein interacting with C-kinase; PIP2, phosphatidylinositol 4,5-bisphosphate; PSD-95, postsynaptic density 95 kDa

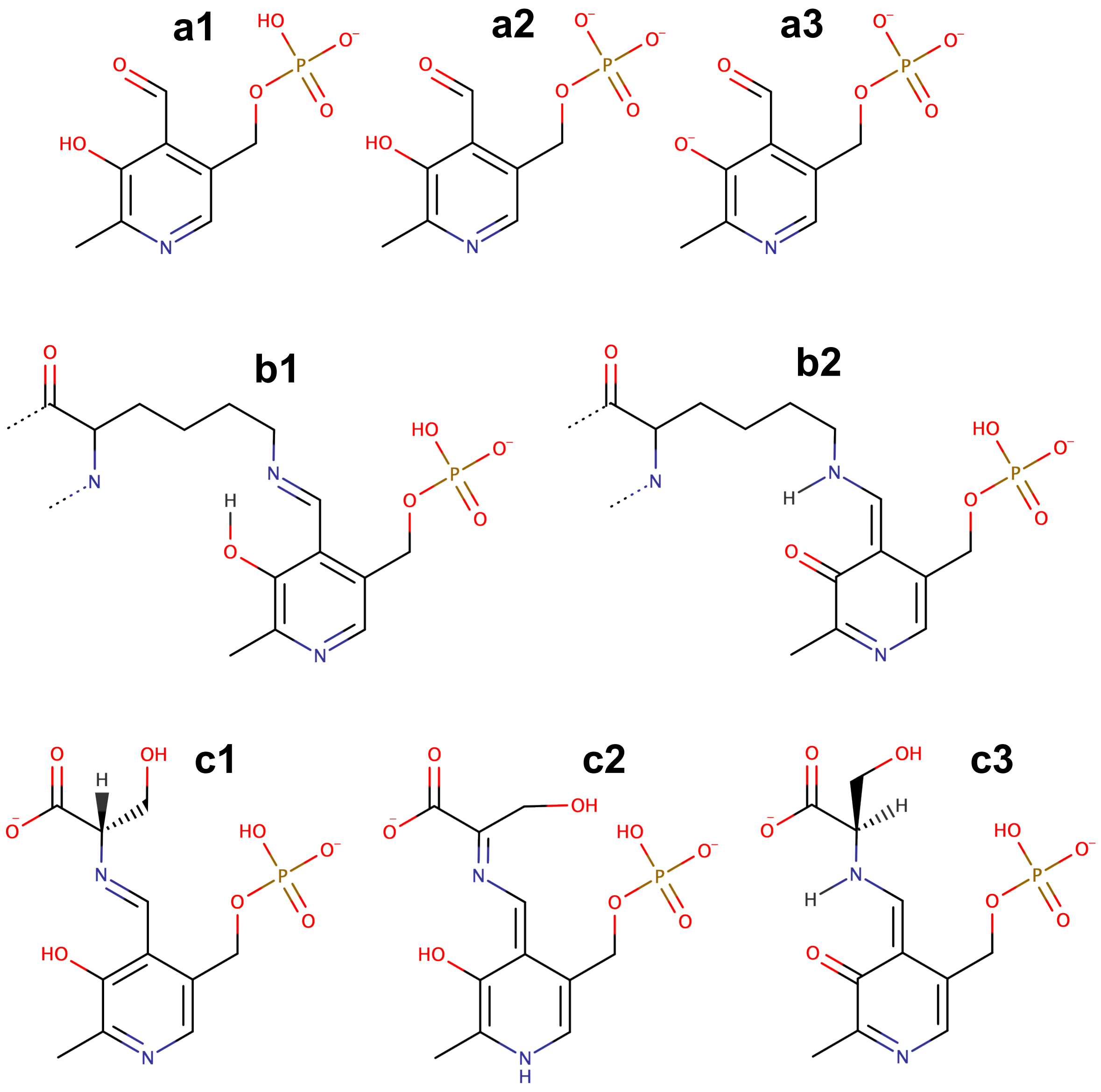

**Supplementary Figure 1**. **Chemical structures of PLP, PLP-Lys56 and PLP-L-Ser, PLP-D-Ser and PLP-planar intermediate.** (**a1, a2, a3**) Chemical structures of three different protonated forms of pyridoxal -5' phosphate (PLP). The minus-1 form is predicted to be most stable below pH 6.8 (**a1**), the minus-2 form between pH 6.8–8.0 (**a2**), and the minus-3 form above pH 8.0 (**a3**). (**b1, b2**) Two tautomers of PLP-Lys56 (internal aldimine). The Schiff base nitrogen not protonated in (**b1**). In both, the aldehyde oxygen has been replaced by a nitrogen and the PLP is now more accurately named a pyridoxamine 5'-phosphate, PMP. (**c1**) External aldimine with L-serine. (**c2**) The 'quinoid' intermediate in which the Cα loses a proton and becomes planar. The Cα can acquire a proton from Ser 84 to give the D-serine adduct (**c3**), or the serine hydroxyl can be protonated and lost as a water. (**c3**) External aldimine with D-serine. MarvinSketch 20.20 (https://www.chemaxon.com) was used to draw chemical structures and predict protonation and tautomeric states.

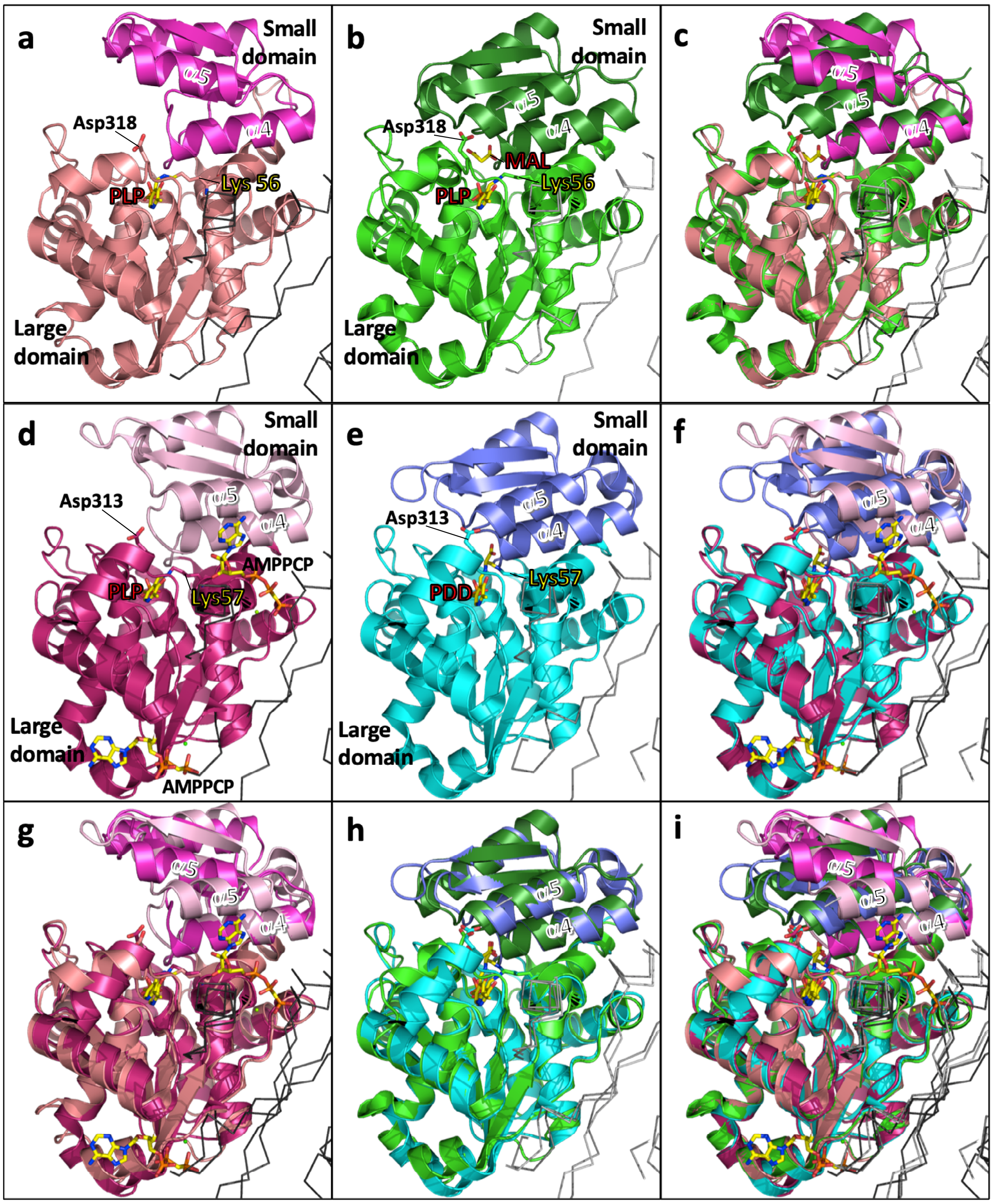

**Supplementary Figure 2. Comparison of 'open' and 'closed' structures of human and yeast SR.**
(**a**) The 'open' 1.89 Å structure of *h*SR (pdb code: 6SLH). The view is from the second subunit in the dimer (Cα ribbon, dark grey). (**b**) The 'closed' 1.5 Å structure of *h*SR with PLP and malonate/MAL (pdb code: 3L6B). The N-terminus of the α4-helix interacts with a malonate carboxylic acid and the N-terminus of the α5-helix interacts with Asp 318 from the conserved SGGNV**D** sequence motif. (**c**) Superposition of (**a**) and (**b**) on large domains. (**d**) The 'open' 1.9 Å yeast structure containing AMPPCP (pdb code: 1WTC). The adenine ring of AMPPCP is close to the α4 and α5 helices. (**e**) The 'closed' 1.7 Å yeast structure (pdb code: 2ZPU) with PDD (N-5'-phosphopyridoxyl)-D-alanine) covalently attached to Lys 57. (**f**) Superposition of (**d**) and (**e**) on large domains. (**g**) Superposition of (**a**) and (**d**) on large domains. Note the relative shift of the α5 helices (**h**) Superposition of (**b**) and (**e**) on large domains. (**i**) Superposition of (**a**), (**b**), (**d**) and (**e**) on large domains.

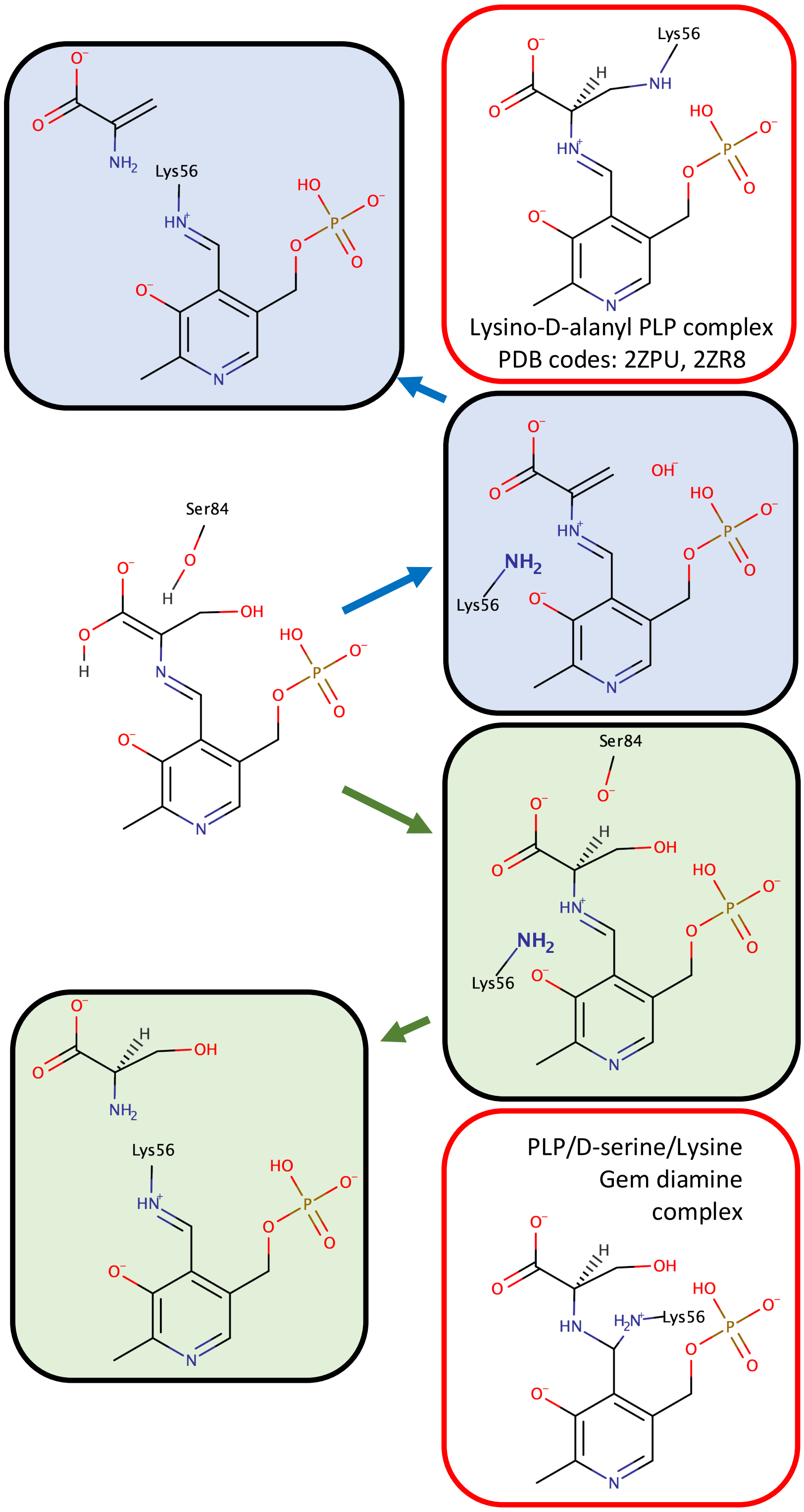

**Supplementary Figure 3**. **Serine racemase chemistry and lysino-d-alanyl and gem-diamine structures**. The reaction products of SR are normally aminoacrylate (blue pathway) and D-serine (green pathway)[33]. At the end of the catalytic cycle the catalytic lysine normally replaces the external aldimine with an internal aldimine. However, with *S. pombe* SR, lysino-D-alaynyl structures were observed and recently with a thermophilic serine hydroxymethyltransferase, a gem diamine structure has been observed (pdb code: 6TI4). MarvinSketch 20.20 (https://www.chemaxon.com) was used to draw chemical structures and predict protonation and tautomeric states.

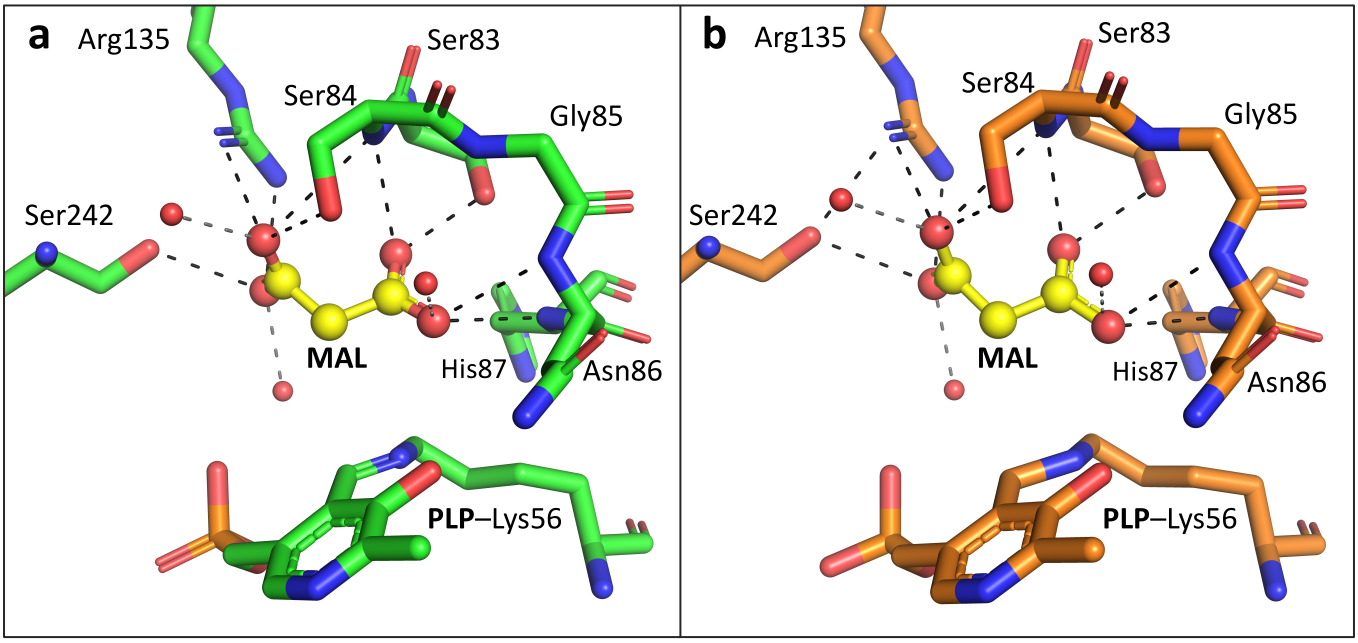

**Supplementary Figure 4. Malonate binding sites in human SR crystal structures (6ZSP and 3L6B).** (**a**) Malonate binding site in 1.6 Å ATP-bound structure (pdb code: 6ZSP). Each of the four oxygens (red atoms) of malonate (yellow carbons) is within H-bonding distance (black dashed lines) of two or three H-bond donors. These include: 5 NHs (blue atoms) from Arg 135 side-chain; main-chain NHs of Ser 84, Asn 86, and His 87; 3 OHs from serine residues 83, 84, and 242; and 3 waters. PLP is covalently attached to Lys 56. (**b**) Malonate binding site of previous SR structure without ATP (pdb code: 3L6B)[3]. The position of malonate and PLP and the organisation of the binding site is virtually identical to that shown in (**a**).

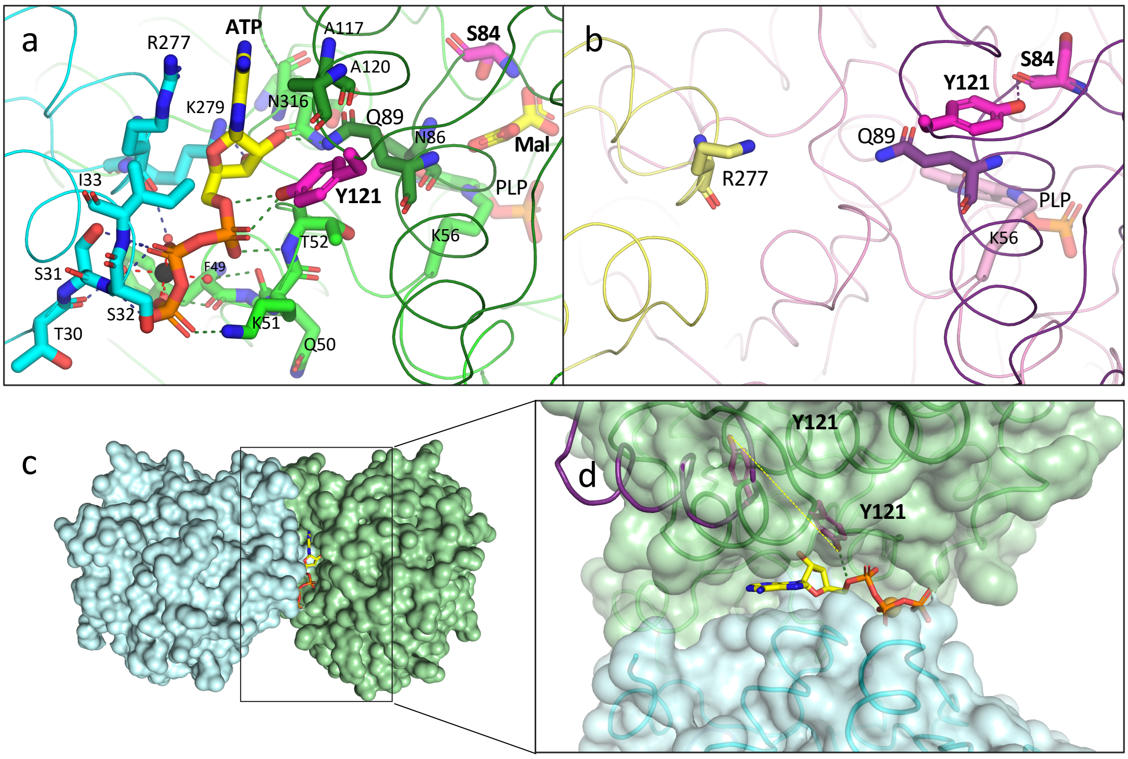

**Supplementary Figure 5. Tyr121 can form hydrogen bonds to ATP or main-chain carbonyl of Ser 84**. (**a**) Close-up of ATP binding site from 1.6 Å ATP/malonate-bound structure (pdb code: 6ZSP) showing ATP (yellow carbons) H-bonded to residues in the small domain (dark green) and large domain (light green) of subunit A in human SR. Tyr121 forms H-bonds to the $\alpha$-phosphate and Gln 89 (side chain in two conformers) forms H-bonds with oxygens from the ribose. Two of the waters (small red spheres) that coordinate the ATP-associated $Mg^{2+}$ ion (black sphere) also form H-bonds with the large domain of the second subunit in the dimer (cyan carbons). (**b**) Equivalent view of new SR holoenzyme structure (pdb code: 6ZUJ). Note the difference in orientation of the side chain of Tyr 121, which is now H-bonded to Ser 84. R277 side-chain is disordered (not shown). (**c**) Surface representation of ATP/malonate-bound structure in similar orientation to (**a**). (**d**) Tyr 121 is oriented towards the ATP site and contributes to the formation of a tightly-fitting ATP binding pocket. The surface is semitransparent, and the position adopted by Tyr 121 at the C-terminal end of the $\alpha$5 helix (PDCKKLAIQAY) in the SR holoenzyme structure is also shown (purple ribbon, tyrosine as stick). The two positions occupied by the terminal hydroxyls of Tyr121 in the two structures are denoted by a dotted yellow line representing a distance of 10 Å.

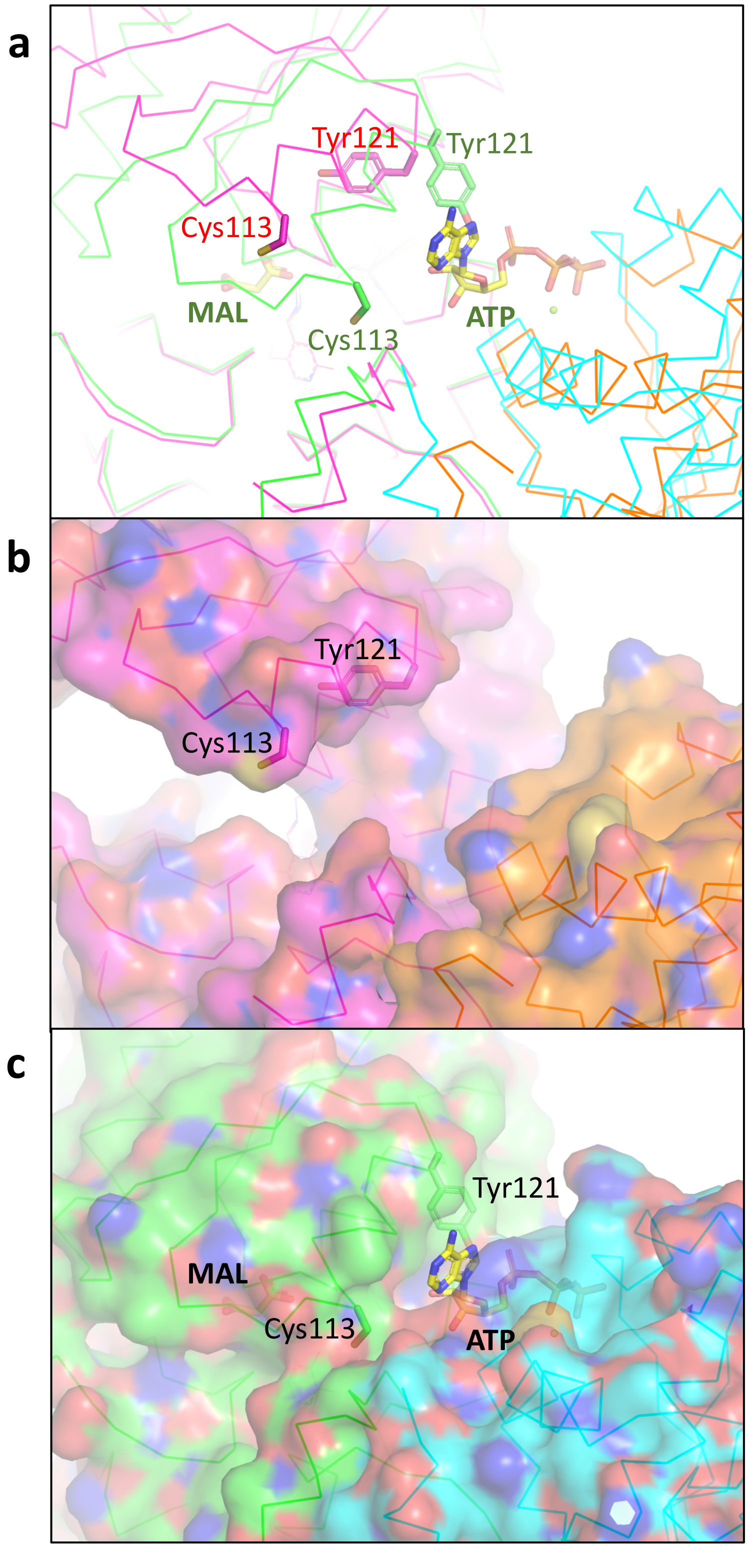

**Supplementary Figure 6. The side-chain of Cys 113 is accessible in the 'open' human SR structure but is buried in the 'closed' structure**. (**a**) Comparison of 'open' and 'closed' structures (Ca traces; ATP, malonate (MAL), and side-chains of Cys 113 and Tyr 121 shown as sticks). (**b**) 'Open' structure as in (a) with semitransparent surface shown – note yellow surface area corresponding to SG of Cys113 (**c**) 'Closed' structure as in (a) with semi-transparent surface shown. The SG of Cys 113 is buried.

```
SRR_HUMAN    ------------MCAQYCISFADVEKAHINIRDSIHLTPVLTSSILNQLTGRNLFFKCELF 49
SRR_MOUSE    ------------MCAQYCISFADVEKAHINIQDSIHLTPVLTSSILNQIAGRNLFFKCELF 49
SRR_RAT      ------------MCAQYCISFADVEKAHLNIQDSVHLTPVLTSSILNQIAGRNLFFKCELF 49
SRR_SCHPO    -----------MSDNLVLPTYDDVASASERIKKFANKTPVLTSSTVNKEFVAEVFFKCENF 50
F5CAQ9_MAIZE MGSKDGTGDISEAQGYAADIDSIREAQARIAPYVHRTPVMSSTSIDAMVGKKLFFKCECF 60
                                  .: .*  .*    : ***::*: ::     ::***** *

SRR_HUMAN    QKTGSFKIRGALNAVRSLVPDALERKPKAVVTHSSGNHGQALTYAAKLEGIPAYIVVPQT 109
SRR_MOUSE    QKTGSFKIRGALNAIRGLIPDTPEEKPKAVVTHSSGNHGQALTYAAKLEGIPAYIVVPQT 109
SRR_RAT      QKTGSFKIRGALNAIRGLIPDTLEGKPKAVVTHSSGNHGQALTYAAKLEGIPAYIVVPQT 109
SRR_SCHPO    QKMGAFKFRGALNALSQLNE---AQRKAGVLTFSSGNHAQAIALSAKILGIPAKIIMPLD 107
F5CAQ9_MAIZE QKAGAFKIRGASNSIFALDD---EQVSKGVVTHSSGNHAAAVALAAKLRGIPAHIVIPRN 117
             ** *:**:*** *::  *          .*:*.*****. *:: :**: **** *::*

SRR_HUMAN    APDCKKLAIQAYGASIVYCEPSDESRENVAKRVTEETEGIMVHPNQEPAVIAGQGTIALE 169
SRR_MOUSE    APNCKKLAIQAYGASIVYCDPSDESREKVTQRIMQETEGILVHPNQEPAVIAGQGTIALE 169
SRR_RAT      APNCKKLAIQAYGASIVYSEPSDESRENVAQRIIQETEGILVHPNQEPAVIAGQGTIALE 169
SRR_SCHPO    APEAKVAATKGYGGQVIMYDRYKDDREKMAKEISEREGLTIIPPYDHPHVLAGQGTAAKE 167
F5CAQ9_MAIZE APASKVENVKCYGGHIIWSDASIESREYVSKRVQEETGAVLIHPINSKYTISGQGTVSLE 177
             ** .*    : **. ::  :   :.** :::.: :.    :: * :   .::**** : *

SRR_HUMAN    VLNQVPLVDALVVPVGGGGMLAGIAITVKALKPSVKVYAAEPSNADDCYQSKLKGKLMPN 229
SRR_MOUSE    VLNQVPLVDALVVPVGGGGMVAGIAITIKALKPSVKVYAAEPSNADDCYQSKLKGELTPN 229
SRR_RAT      VLNQVPLVDALVVPVGGGGMVAGIAITIKTLKPSVKVYAAEPSNADDCYQSKLKGELTPN 229
SRR_SCHPO    LFEEVGPLDALFVCLGGGGLLSGSALAARHFAPNCEVYGVEPEAGNDGQQSFRKGSIVH- 226
F5CAQ9_MAIZE LLEQVPEIDTIIVPISGGGLISGVALAAKAINPSIRILAAEPKGADDSAQSKAAGKIIT- 236
             ::::*  :*::.* :.***:::* *:: : : *. .: ..**. .:*  **   *.:

SRR_HUMAN    LYPPETIADGVK-SSIGLNTWPIIRDLVDDIFTVTEDEIKCATQLVWERMKLLIEPTAGV 288
SRR_MOUSE    LHPPETIADGVK-SSIGLNTWPIIRDLVDDVFTVTEDEIKYATQLVWGRMKLLIEPTAGV 288
SRR_RAT      LHPPETIADGVK-SSIGLNTWPIIRDLVDDVFTVTEDEIKYATQLVWERMKLLIEPTAGV 288
SRR_SCHPO    IDTPKTIADGAQTQHLGNYTFSIIKEKVDDILTVSDEELIDCLKFYAARMKIVVEPTGCL 286
F5CAQ9_MAIZE LPSTNTIADGLR-AFLGDLTWPVVRDLVDDVIVVDDTAIVDAMKMCYEILKVAVEPSGAI 295
             :   :***** :   :*  *: :::: ***::.* :  :  . ::    :*: :**:. :

SRR_HUMAN    GVAAVLSQHFQTVSP--EVKNICIVLSGGNVDLTSSITWVKQAERPASYQSVSV 340
SRR_MOUSE    ALAAVLSQHFQTVSP--EVKNVCIVLSGGNVDLT-SLNWVGQAERPAPYQTVSV 339
SRR_RAT      GLAAVLSQHFQTVSP--EVKNICIVLSGGNVDLT-SLSWVKQAERPAP------ 333
SRR_SCHPO    SFAAARAMKE-----KLKNKRIGIIISGGNVDIERYAHFLSQ------------ 323
F5CAQ9_MAIZE GLAAALSDEFKQSSAWHESSKIGIIVSGGNVDLGTL--WQSMYKH--------- 338
             ..**. : .        : ..: *::*******:    :
```

**Supplementary Figure 7. Serine racemase sequence alignment**. SR amino acid sequences from *H. sapiens* (SRR_HUMAN), *M. musculus* (SRR_MOUSE), *R. norvegicus* (SRR_RAT), *S. pombe* (SRR_SCHPO) and *Z. mays* (F5CAQ9_MAIZE) were aligned with ClustalO using sequences from UniProt[34]. The small domain of *h*SR (residues 78–155) is underlined. Residues (S84, G88, Q89, T92, I104, A117, I118) that are within 3.8 Å of the side chain of Tyr 121 (Y121) in the 'open' structure (pdb code: 6ZUJ) are highlighted in red on the *h*SR (SRR_HUMAN) sequence. Note that G88 and T92 are alanines in the *S. pombe* and *Z. Mays* sequences. Conserved residues are indicated by * underneath alignment. Highlighted residues are C113 (the nitrosylation site in SRR_MOUSE[28]) and T71 and T227 (the phosphorylation sites in SRR_MOUSE[35]).

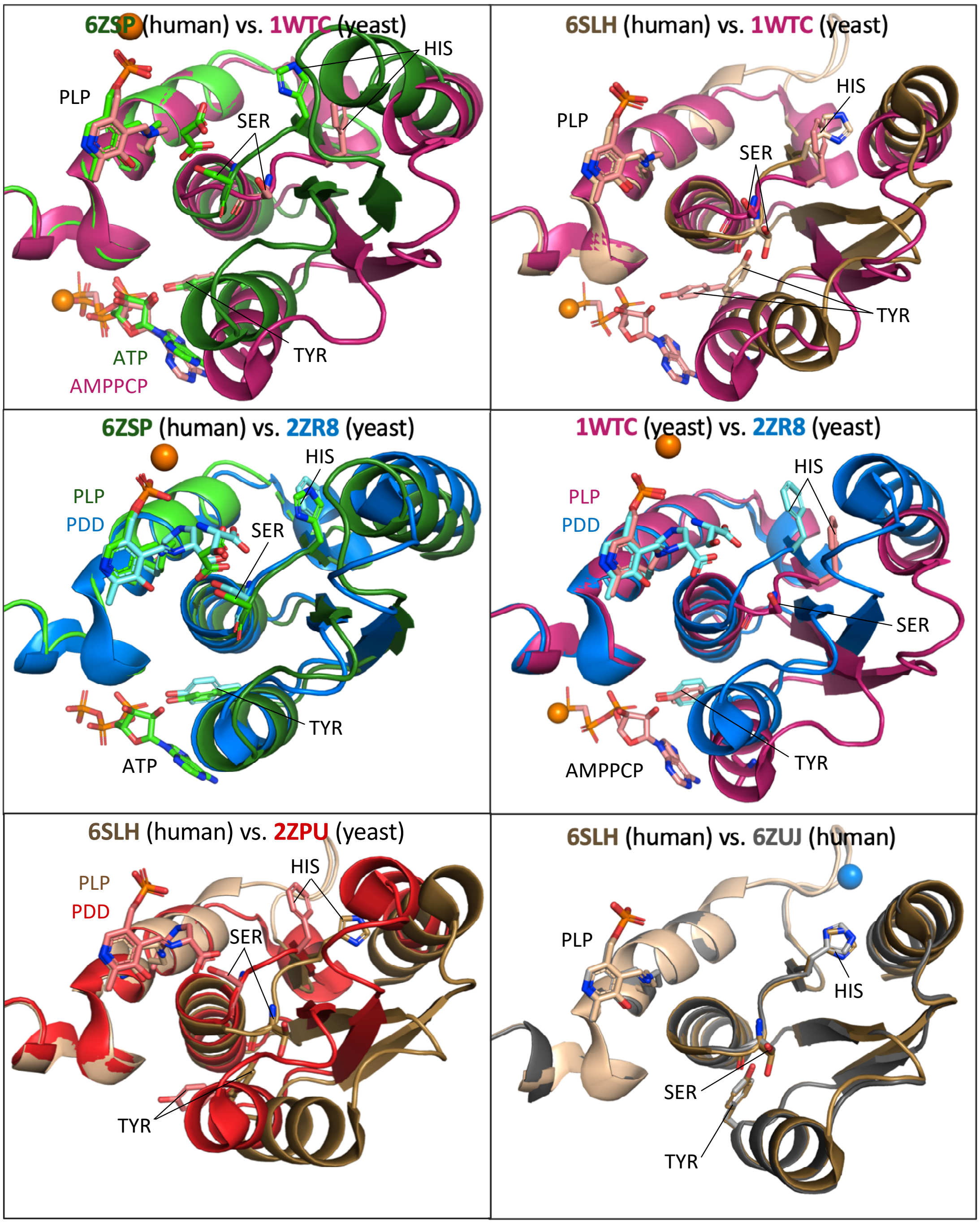

**Supplementary Figure 8. Superpositions of human *holo* and ATP/malonate-bound SR with yeast orthologues**. The displayed structures are: **6ZSP** (green; human malonate-bound with ATP); **1WTC** (magenta; yeast *holo* with ATP analogue, AMPPCP); **6SLH** (small domain brown, side-chains + large domain light pink; human *holo*); **2ZR8** (blue; with both L-Ser and modified PLP – called PDD see supplementary Table 5); **2ZPU** (red; yeast *holo*); **6ZUJ** (grey, new XChem human *holo*). The residues in stick are TYR (Tyr 121 in human, Tyr 119 in yeast), SER (Ser 84 in human, Ser 82 in yeast) and HIS (His 82 in human, His 80 in yeast).

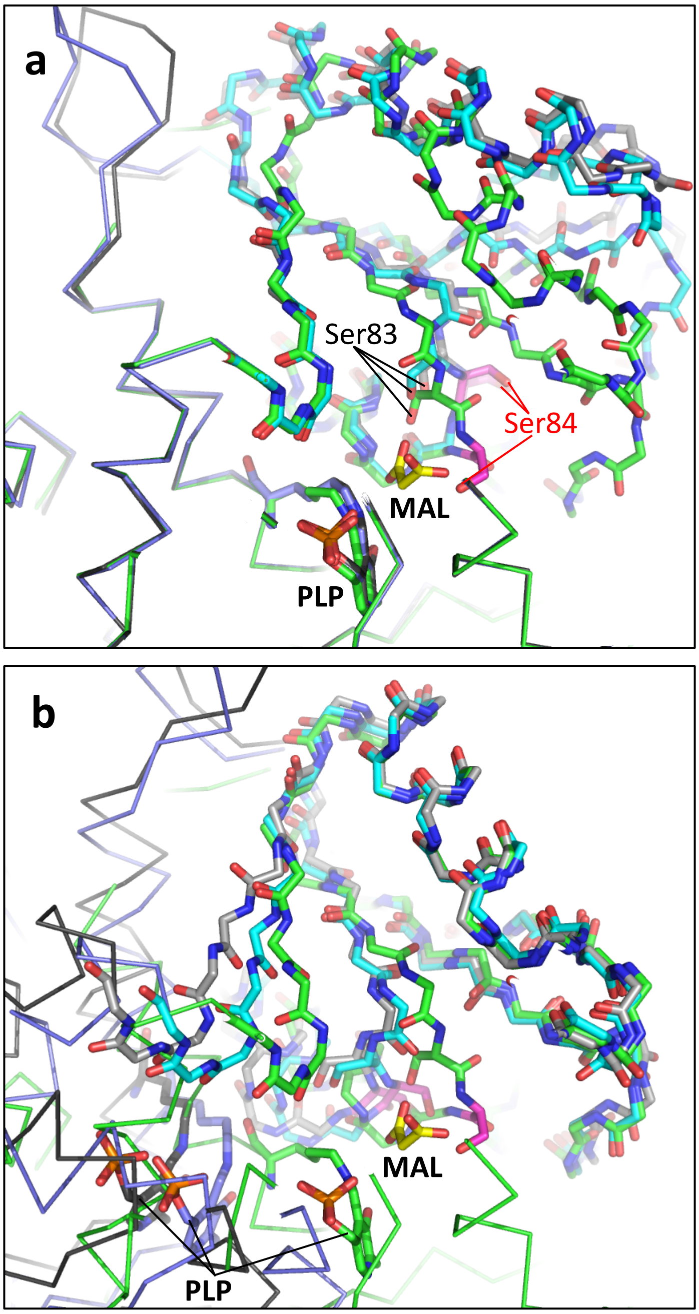

**Supplementary Figure 9. Superpositions of ‘closed’ SR ATP/malonate-bound structure with ‘open’ SR subunits A and C.** (**a**) The ‘closed’ *h*SR (green carbons) complex with malonate/MAL (stick; yellow carbons) and ATP (not visible) is shown with large domain as Cα ribbon and small domain as backbone sticks. The side-chain of Ser 84 (stick magenta carbons) is close to the malonate. The PLP (stick; green carbons) is covalently attached to Lys 56. Superposed on Mal/ATP large domain are the A subunit of an XChem structure (blue/cyan carbons, compound **13**-x0430 – compound not observed in A subunit) and the C subunit of an XChem structure (grey carbons, compound **1**-x0495 - compound not seen in C subunit). Side-chain hydroxyls of Ser 83 and 84 are arrowed. (**b**) The closed *h*SR (green carbons) complex with malonate/MAL (stick; yellow carbons) and ATP (not visible) as in (a). The XChem A and C subunits are superposed based on residues 78–81 and 101–148 of the small domain (residues 77–155). Note large relative shifts of PLP and large domain.

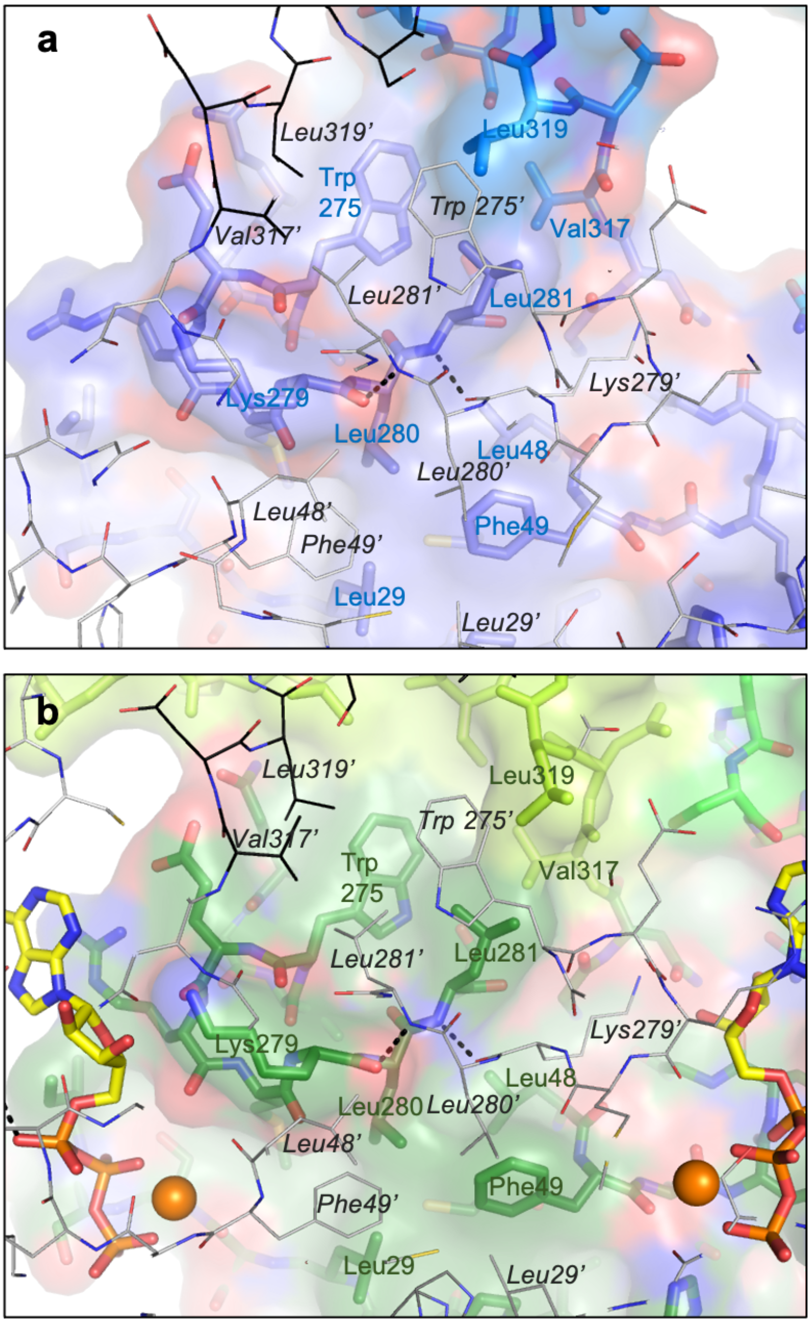

**Supplementary Figure 10. The hydrophobic core of the dimer interface in 'open' and 'closed' hSR structures.** (**a**) The AC dimer interface from an 'open' XChem structure (pdb code: 6ZUJ). The A subunit (stick; blue carbons) is shown with a semitransparent surface. The C subunit (line representation; grey carbons) is nearer the viewer. Residues are labelled with blue letters for the A subunit and black letters for the C subunit. Intersubunit hydrogen bonds between CO of Lys 279 to the NH of Leu 281' and from the NH of Leu 281 to CO of Lys 279' are indicated by dotted lines. (**b**) The AB dimer interface from the 'closed' malonate/ATP structure (pdb code: 6ZSP). The A subunit (stick; green carbons) is shown with a semitransparent surface. The B subunit (line representation; grey carbons) is nearer the viewer. ATPs (stick, yellow carbons) and magnesium ions (orange spheres) are also present.

```
SRR_HUMAN     ------------MCAQYCISFADVEKAHINIRDSIHLTPVLTSSILNQLTGRNLFFKCELF 49
SRR_RAT       ------------MCAQYCISFADVEKAHLNIQDSVHLTPVLTSSILNQIAGRNLFFKCELF 49
SRR_SCHPO     -----------MSDNLVLPTYDDVASASERIKKFANKTPVLTSSTVNKEFVAEVFFKCENF 50
F5CAQ9_MAIZE  MGSKDGTGDISEAQGYAADIDSIREAQARIAPYVHRTPVMSSTSIDAMVGKKLFFKCECF 60
SDHL_HUMAN    --------------------------MMSGEPLHVKTPIRDSMALSKMAGTSVYLKMDSA 34
SDHL_RAT      --------------------------MAAQESLHVKTPLRDSMALSKVAGTSVFLKMDSS 34
                                            **:   *   :.      .:::* :

SRR_HUMAN     QKTGSFKIRGALNAVRSLVPDALERKPKAVVTHSSGNHGQALTYAAKLEGIPAYIVVPQT 109
SRR_RAT       QKTGSFKIRGALNAIRGLIPDTLEGKPKAVVTHSSGNHGQALTYAAKLEGIPAYIVVPQT 109
SRR_SCHPO     QKMGAFKFRGALNALSQLNE---AQRKAGVLTFSSGNHAQAIALSAKILGIPAKIIMPLD 107
F5CAQ9_MAIZE  QKAGAFKIRGASNSIFALDD---EQVSKGVVTHSSGNHAAAVALAAKLRGIPAHIVIPRN 117
SDHL_HUMAN    QPSGSFKIRGIGHFCKRW----AKQGCAHFVCSSAGNAGMAAAYAARQLGVPATIVVPST 90
SDHL_RAT      QPSGSFKIRGIGHLCKMK----AKQGCKHFVCSSAGNAGMATAYAARRLGLPATIVVPST 90
              *  *:**:**   :                 .:   *:** . * : :*:  *:** *::*

SRR_HUMAN     APDCKKLAIQAYGASIVYCEPSDESRENVAKRV-TEETEGIMVHPNQEPAVIAGQGTIAL 168
SRR_RAT       APNCKKLAIQAYGASIVYSEPSDESRENVAQRI-IQETEGILVHPNQEPAVIAGQGTIAL 168
SRR_SCHPO     APEAKVAATKGYGGQVIMYDRYKDDREKMAKEI-SEREGLTIIPPYDHPHVLAGQGTAAK 166
F5CAQ9_MAIZE  APASKVENVKCYGGHIIWSDASIESREYVSKRV-QEETGAVLIHPINSKYTISGQGTVSL 176
SDHL_HUMAN    TPALTIERLKNEGATVKVVGELLDEAFELAKALAKNNPGWVYIPPFDDPLIWEGHASIVK 150
SDHL_RAT      TPALTIERLKNEGATVEVVGEMLDEAIQLAKALEKNNPGWVYISPFDDPLIWEGHTSLVK 150
              :*   .    :  *. :       :.   ::: :  :.     : * :      *:  :

SRR_HUMAN     EVLNQVPL-VDALVVPVGGGGMLAGIAITVKALK-PSVKVYAAEPSNADDCYQSKLKGKL 226
SRR_RAT       EVLNQVPL-VDALVVPVGGGGMVAGIAITIKTLK-PSVKVYAAEPSNADDCYQSKLKGEL 226
SRR_SCHPO     ELFEEVGP-LDALFVCLGGGGLLSGSALAARHFA-PNCEVYGVEPEAGNDGQQSFRKGSI 224
F5CAQ9_MAIZE  ELLEQVPE-IDTIIVPISGGGLISGVALAAKAIN-PSIRILAAEPKGADDSAQSKAAGKI 234
SDHL_HUMAN    ELKETLWEKPGAIALSVGGGGLLCGVVQGLQEVGWGDVPVIAMETFGAHSFHAATTAGKL 210
SDHL_RAT      ELKETLSAKPGAIVLSVGGGGLLCGVVQGLREVGWEDVPIIAMETFGAHSFHAAVKEGKL 210
              *: : :     .:: : :.***::.*  .    : .    .  : . *    ...    :    *.:

SRR_HUMAN     MPNLYPPETIADGVK-SSIGLNTWPIIRDLVDDIFTVTEDEIKCATQLVWERMKLLIEPT 285
SRR_RAT       TPNLHPPETIADGVK-SSIGLNTWPIIRDLVDDVFTVTEDEIKYATQLVWERMKLLIEPT 285
SRR_SCHPO     VH-IDTPKTIADGAQTQHLGNYTFSIIKEKVDDILTVSDEELIDCLKFYAARMKIVVEPT 283
F5CAQ9_MAIZE  IT-LPSTNTIADGLR-AFLGDLTWPVVRDLVDDVIVVDDTAIVDAMKMCYEILKVAVEPS 292
SDHL_HUMAN    VS-LPKITSVAKALGVKTVGAQALKLFQEHPIFSEVISDQEAVAAIEKFVDDEKILVEPA 269
SDHL_RAT      VT-LPKITSVAKALGVNTVGAQTLKLFYEHPIFSEVISDQEAVTAIEKFVDDEKILVEPA 269
                 :    ::*..     :*   :   :. :        .: :      . :       *: :**:

SRR_HUMAN     AGVGVAAVLSQHFQTVSP-------EVKNICIVLSGGNVDLTSSITWVKQAERPASYQSVSV 340
SRR_RAT       AGVGLAAVLSQHFQTVSP-------EVKNICIVLSGGNVDLT-SLSWVKQAERPAP------ 333
SRR_SCHPO     GCLSFAAARAMKE-----KL-----KNKRIGIIISGGNVDIERYAHFLSQ------------ 323
F5CAQ9_MAIZE  GAIGLAAALSDEFKQSSAWH-----ESSKIGIIVSGGNVDLGTL--WQSMYKH--------- 338
SDHL_HUMAN    CGAALAAVYSHVIQKLQLEGNLRTPLPSLVVIVCGGSNISLAQLRALKEQLGMTNRLPK--- 328
SDHL_RAT      CGAALAAVYSGVVCRLQAEGRLQTPLASLVVIVCGGSNISLAQLQALKAQLGLNELLK---- 327
                 ..**. :                  . : *: .*.*:.:
```

**Supplementary Figure 11. Serine dehydratase and serine racemase sequence alignment.** SR amino acid sequences are as described in Supplementary Fig. 7. Structures for all sequences are available in the PDB (see Supplementary Table 5). Serine dehydratase (SDH) sequences for *H. sapiens* (SDHL_HUMAN) and *R. norvegicus* (SDHL_RAT) (isoform 2) were obtained from UniProt[34]. Conserved residues are indicated by * underneath alignment. Residues highlighted in red on the *h*SR sequence (T52, K56, H82, S84, Q89, Y121, R135, E136). Note the conservation of S83, G85, and N86.

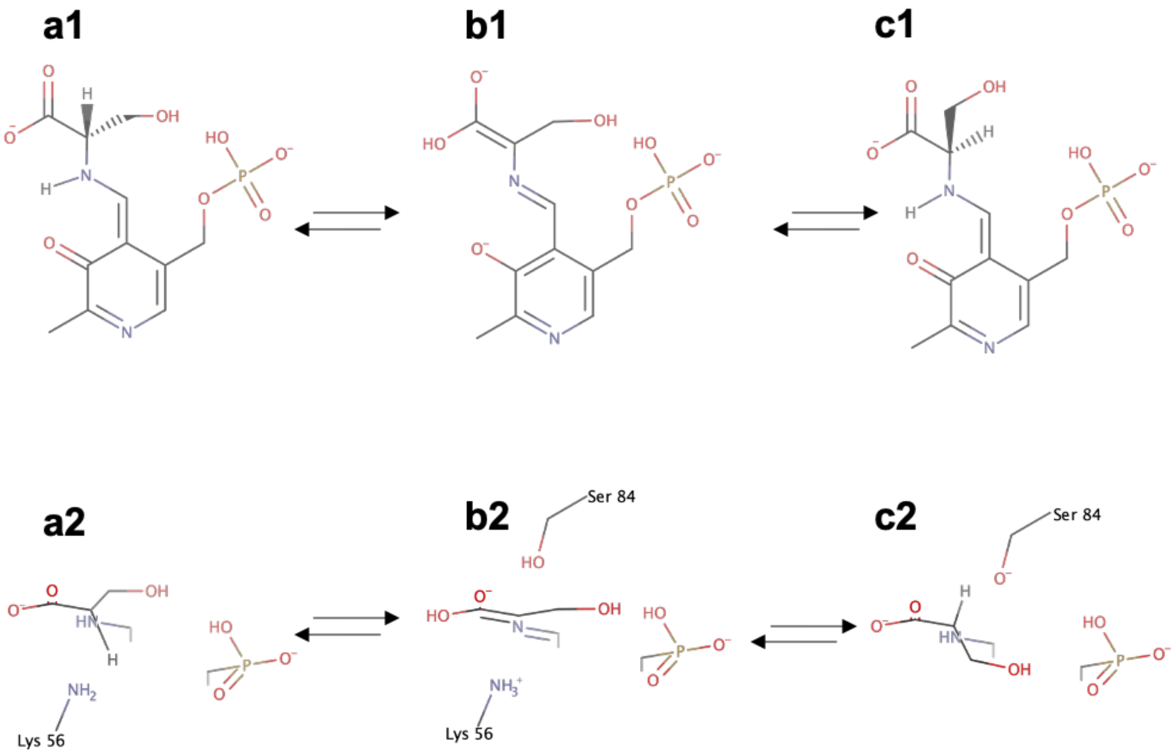

**Supplementary Figure 12. Roles of lysine 56 and serine 84 in proton exchange in human SR**. (**a1, a2**) Orthogonal view of an L-serine PLP external aldimine. The view in (**a2**) is looking down from above (**a1**). Only the uppermost atoms of (**a1**) are shown in (**a2**). Lys 56 extracts the Cα proton between (**a1/a2**) and (**b1/b2**) or donates the proton in the reverse process. (**b1/b2**) Orthogonal views of a planar intermediate form in which the Cα of serine has no hydrogen. (**c1/c2**) Orthogonal view of a D-serine PLP external aldimine. Ser 84 donates (**b** to **c**) or extracts (**c** to **b**) the Cα proton. MarvinSketch 20.20 (https://www.chemaxon.com) was used to draw chemical structures and predict protonation and tautomeric states.

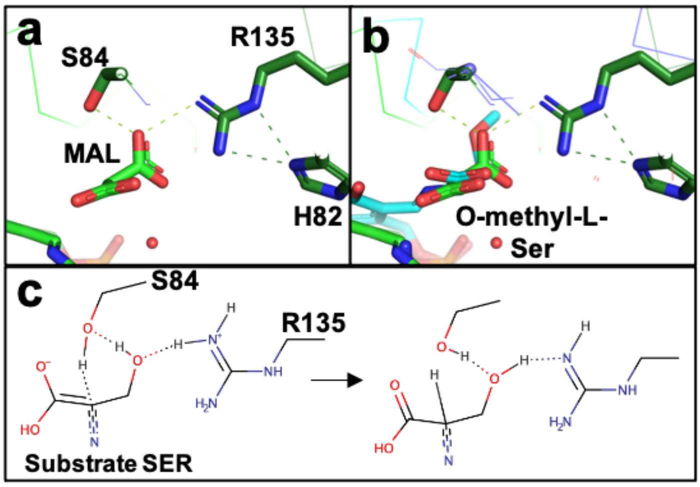

**Supplementary Figure 13**. Does Arginine 135 protonate Ser 84 via the side-chain of the substrate serine? (**a**) A hydrogen bond network between Ser 84, malonate (MAL), Arg 135 and His 82 in the 1.6 Å hSR structure (pdb code: 6ZSP). (**b**) The structure of SDH in complex with O-methyl-serine (pdb code: 1PWH) is shown superposed on the 1.6 Å malonate/ATP structure. (**c**) Proposed mechanism demonstrating how Arg 135 could protonate the hydroxyl of the substrate serine as it in turn protonates Ser 84. The mechanism suggests that Arg 135 could protonate the hydroxyl of the substrate serine; the substrate serine protonates Ser 84, which in turn protonates the Cα of the substrate. Arg 135 is conserved in serine racemase sequences but is not present in serine dehydratases (see Supplementary Fig. 11).

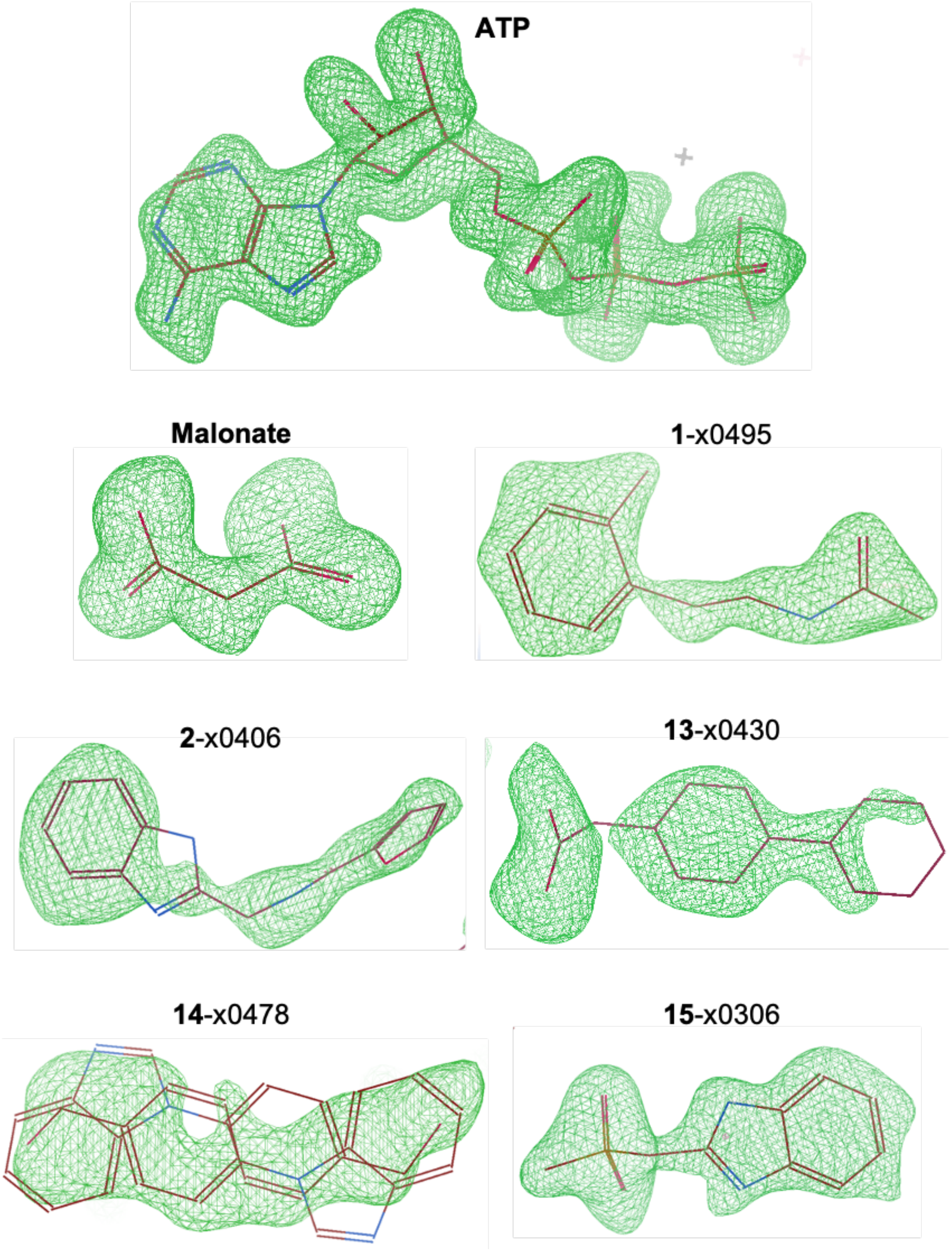

**Supplementary Figure 14. OMIT maps of ATP, malonate, and bound XChem ligands**. Fo-Fc electron density (green mesh) contoured to 3 σ showing evidence for bound ligand. OMIT maps were generated by removing the ligand from the structure and running five cycles of refinement in REFMAC. Shown here are ATP and malonate (pdb code: 6ZSP), **1**-x0495 (7NBG), **2**-x0406 (7NBH), **13**-x0430 (7NBC), **14**-x0478 (7NBD), and **15**-x0306 (7NBF). Compound **14**-x0478 sits on the twofold axis of the dimer and observed electron density is due to two (2-fold related) equivalent binding modes. Occupancies range from 0.7–1.0 (2 x 0.5 for **14**-x0478).

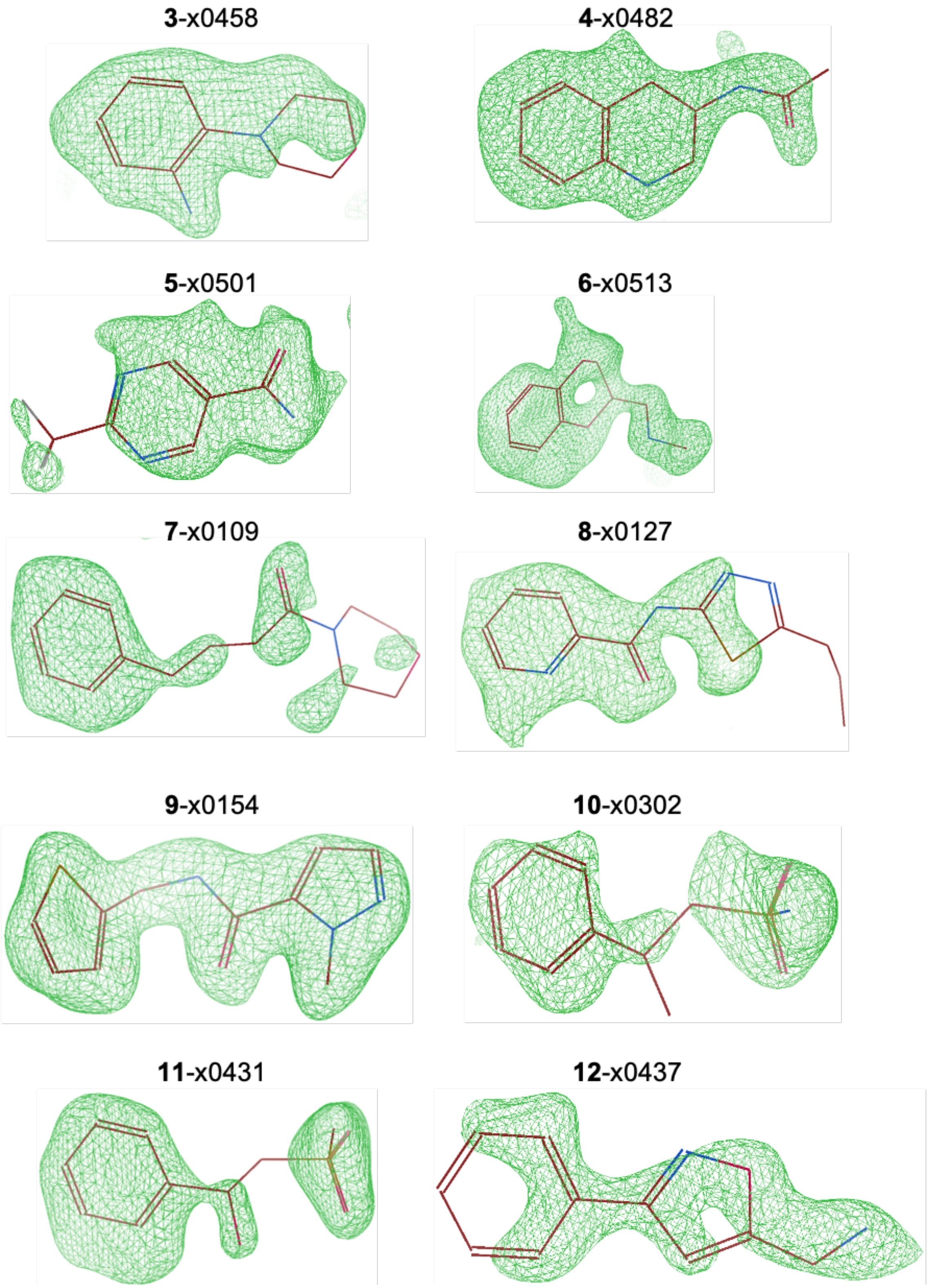

**Supplementary Figure 15. OMIT maps of XChem ligands bound to SR**. Fo-Fc electron density (green mesh) contoured to 3 σ showing evidence for bound XChem ligand. OMIT maps were generated by removing the ligand from the structure and running five cycles of refinement in REFMAC. Occupancies range from 0.7–1.0.

**Supplementary Discussion.**

**The evolution and regulation of the activity of human serine racemase in the CNS.**

In the CNS, SR can localise to specific regions in cells and its activity is tightly spatially and temporally regulated by a number of protein interactions[24-27,36-41] and post-translational modifications[28,35,36,42,43]. The emerging model suggests that astrocytes normally synthesize L-serine, which is then transported to neurons where D-serine is synthesised and utilized at synapses[24,40,44,45]. However, in the diseased or damaged brain, astrocytes can synthesize D-serine and the extra-synaptic activity of D-serine is neurotoxic[45]. SR interacts with postsynaptic proteins such as PICK1[26,46], GRIP[25,47], and PSD-95[41] to increase production of D-serine through unspecified structural mechanisms (see Supplementary Table 6 for a list of physiological modulators of SR). SR ubiquitination and degradation is tempered by its interaction with Golga3[27] and DISC1[32,40,48,49], the latter of which is linked to diminished D-serine and a schizophrenic phenotype when mutated. Understanding how SR activity is regulated in cells may facilitate optimisation of therapeutic inhibitors and activators of SR. While it is clear that serine racemase and NMDA receptors play important roles in animals with a central nervous system, from *C. elegans* to man[50], more research remains to be done to understand exactly how SR activity is regulated *in vivo*.

How serine racemase activity evolved is also an intriguing question, one possibility is that SR evolved from a progenitor protein that had serine dehydratase activity but lacked racemisation activity. Human serine dehydratase (SDH), the closest human homologue of hSR with which it shares a common ancestor, is about 75-fold more active at elimination than human SR[9]. However, human SDH has no detectable elimination activity against D-serine. Racemisation is believed to have emerged as a side-reaction of the elimination function[51]. Comparison of structures of rat SDH with hSR (Fig. 8) suggested that Ser 84 is the crucial catalytic residue for racemisation[3], and this was confirmed by site-directed mutagenesis experiments in which the activity of a S84A mutant was determined[9]. The hSR S84A mutant had no detectable racemase activity against L-serine or D-serine. The S84A mutant also had no detectable elimination activity against D-serine while its activity against L-serine was comparable to wild-type[9]. However, the A65S mutant of SDH (Supplementary Fig. 11 for sequence alignment) had no detectable racemase activity[9] but it did acquire $\beta$-elimination activity against D-serine. Wild-type SDH only efficiently catalyses one reaction (the breakdown of L-serine into pyruvate and ammonia) of the four reactions on L- or D-serine catalysed by SR (L to D-Ser, D to L-Ser, L-Ser to pyruvate and ammonia, D-Ser to pyruvate and ammonia). The mutation A65S in SDH gives the enzyme some ability to catalyse the breakdown of D-serine to pyruvate and ammonia[9], supporting the generally accepted roles of Ser 84 and Lys 56 in hSR as the Si and Re-face acid/bases (Supplementary Fig. 12). However, if SR did indeed evolve from a progenitor of SDH, the acquisition of racemisation activity seems to require more than a simple acquisition of a serine at position 84 (=65 in rSD; Fig. 8).

In yeast *S. pombe* SR (*sp*SR), L-serine racemisation activity is over 20-fold lower than its elimination activity[5]. Mouse and human SR have comparable efficiencies for both reactions, with L-serine racemisation efficiency increased by over 10-fold relative to yeast SR while L-serine elimination is mostly unchanged[11,52]. Further, the *in vitro* rate of human SR racemisation is about 100-fold lower than that of alanine racemase, an archetypical amino acid racemase in bacteria[53], which supports the theory racemisation evolved later as a result of catalytic promiscuity[54]. Note, however, that *in vivo* activity is regulated by various interactions (see Supplementary Table 6) and could differ appreciably from *in vitro* activity.

The roles of Ser 84 and Lys 56 in abstracting or donating a proton from the central C$\alpha$ to convert L- to D-serine (or vice versa) is quite well understood (Supplementary Fig. 12). However, two alternate mechanisms have been proposed for the origin of the hydrogen in the $\beta$-elimination reaction. Smith *et al.*[3] state that the elimination reaction requires removal of water with the hydroxyl group from the C$\beta$ of L-serine and hydrogen from the lysyl-$NH_3$ as leaving groups. Another mechanism proposed for SDH states that the phosphate group of PLP acts as a general acid to donate a proton to the leaving hydroxyl group of the serine[7]. A proposed elimination mechanism (Fig. 9d) has the phosphate indirectly protonating the leavinh hydroxyl, via a water. Unexpectedly in a recent neutron study of a PLP-dependent enzyme[55] a deuterium was observed equidistant between the Schiff base and the C-terminal carboxylate of the substrate. How protons move in the catalytic cycles of PLP dependent enzymes is not fully understood. The structure of yeast SR with a lysino-D-alanyl residue (Fig. 8)[5] showed that the catalytic lysine can become covalently attached to the substrate C$\beta$ side-chain[5]. Yamauchi et al.[5] reported that incubation of mouse SR with L-serine caused an enzyme to form with a higher mass than the native enzyme, which may indicate the formation of the lysine-substrate complex even in mammalian SRs. The broad range of possible protonation and tautomeric states[56] of the substrate/PLP conjugate (Supplementary Fig. 1) suggests that further studies to experimentally determine hydrogen positions as the enzyme moves through the racemisation or elimination catalytic cycle would be of interest. Close inspection of our 1.6 Å malonate/ATP structure suggests that the acidity of Ser 84 may be enhanced by Arg 135 (Supplementary Fig. 13), consistent with the conservation of Arg 135 in serine racemase but not serine dehydratase (Supplementary Fig. 11). However, in 2017, Nelson et al. (36) proposed that Lys 114 might play an important role in enhancing the ability of Ser 84 to protonate the substrate and thus produce D-Serine. The proposed mechanism (Fig. 9c) adequately explains the protonation of a planar quinoid, although other possible mechanisms exist (e.g. Supplementary Fig. 13). However, neither mechanism (Fig. 9c, Supplementary Fig. 13) provides a ready explanation for the role of Ser 84 in abstracting the proton from the Ca of a D-serine substrate (Supplementary Fig. 12).

The sensitivity of SR to ATP activation varies among the SR orthologues with a conserved tyrosine. In slime mould (*D. discoideum*) SR, Mg-ATP increases racemisation by 2.5-fold and elimination by 6-fold, indicating the presence of an ATP site[57]. Yeast SR is minimally sensitive to ATP as both reactions increase by less than 2-fold[4]. In human SR, L-serine elimination is most profoundly affected, increasing by 31-fold upon addition of ATP, followed by D-serine elimination which increases by 4-fold and finally L-serine racemisation by almost 2-fold[58,59]. Thus *in vitro*, the L-serine elimination reaction is several times more efficient than the L-serine racemisation reaction[52,58,59]. SDH does not have a conserved tyrosine or an ATP site and is not regulated by nucleotides[9], suggesting ATP binding is coupled to SR's evolutionary divergence from dehydratases.

The role of the Tyr 121 flip appears to be most relevant for the mammalian enzyme, where the complexity of the nervous system necessitates stricter maintenance of D-serine homeostasis in neurons. SR is thought to alternately synthesise and degrade D-serine in brain regions lacking D-amino acid oxidase (DAO), the main degradative enzyme for D-amino acids[60] that is restricted to the cerebellum and brainstem. It has been reported that enhanced astrocytic D-serine synthesis underlies synaptic damage after traumatic brain injury[61] and that neurotoxic astrocytes synthesize SR in Alzheimer's disease[62]. SR mutants deficient in elimination activity accumulate D-serine[63], therefore the elimination function of mammalian SR is critical with respect to preventing D-serine related neuropathologies. Several small molecules and post-translational modifications reduce SR activity in an ATP-dependent manner, including NADH[30], the membrane-localised inhibitor PIP2[24], and S-nitrosylation[28]. The Tyr 121 flip may serve to regulate serine metabolism by preventing basal degradation or synthesis of D-serine when ATP has been displaced by external regulators. When bound, the position of ATP appears to restrict how the small domain moves relative to the large domain, which may explain why ATP enhances racemisation less potently than L-serine elimination.
